# Role of the small protein Mco6 in the mitochondrial sorting and assembly machinery

**DOI:** 10.1101/2023.02.03.527057

**Authors:** Jon V. Busto, Hannah Mathar, Conny Steiert, Eva F. Schneider, Sebastian P. Straub, Lars Ellenrieder, Jiyao Song, Sebastian B. Stiller, Philipp Lübbert, Bernard Guiard, Fabian den Brave, Uwe Schulte, Bernd Fakler, Thomas Becker, Nils Wiedemann

## Abstract

The majority of mitochondrial precursor proteins are imported through the Tom40 β-barrel channel of the translocase of the outer membrane (TOM). The sorting and assembly machinery (SAM) is essential for β-barrel membrane protein insertion into the outer membrane and thus required for the assembly of the TOM complex. Here we demonstrate that the a-helical outer membrane protein Mco6 forms a complex with the mitochondrial distribution and morphology protein Mdm10 as part of the SAM machinery. Moreover, Mco6 also interacts with the subunit Mim1 of the mitochondrial import complex (MIM), which is itself required for the biogenesis of a-helical outer membrane proteins. *MCO6* and *MDM10* display a negative genetic interaction and a *MCO6-MDM10* yeast double mutant contains reduced levels of TOM complex. Cells lacking Mco6 affect the levels of Mdm10 and MIM-subunits associated with assembly defects of the TOM complex. Thus, this work reveals a role of the SAM^Mco6^ complex for the biogenesis of the mitochondrial outer membrane.

## INTRODUCTION

Mitochondria are essential cell organelles of eukaryotic cells required for cellular respiration and the synthesis of Fe-S proteins.^1^ Mitochondrial dysfunction causes diseases affecting the nervous system, muscles and other organs with high energy demand like the liver.^2^ Mitochondria contain β-barrel membrane proteins which form pores in the outer membrane. The porin/VDAC (voltage dependent anion channel) β-barrel channels are required for the exchange of metabolites between the cytosol and the intermembrane space and the β-barrel core subunit Tom40 of the translocase of the outer mitochondrial membrane (TOM) is required for the import of the vast majority of ~1000 nuclear-encoded mitochondrial proteins.^3–6^ All mitochondrial β-barrel proteins are encoded in the nucleus and are imported across the outer membrane by the TOM complex. TIM transfer chaperones in the intermembrane space guide the β-barrel membrane protein precursors through the aqueous environment to the sorting and assembly machinery (SAM/TOB), which is essential for their insertion into the outer mitochondrial membrane.^7,8^

The SAM machinery contains four subunits Sam50 (Tob55/Omp85), Sam35 (Tob38), Sam37 and Mdm10.^9–13^ The C-terminal β-strand of the β-barrel precursor proteins contains a β-signal, which is required to initiate barrel assembly at the SAM complex.^14^ Sam50, the essential core subunit of the SAM machinery, is a β-barrel protein itself and forms a lateral gate between its first and last β-strands.^15^ Sam50 forms a dimer in the outer membrane with the two lateral gates facing each other. Sam35 and Sam37 are peripheral membrane proteins which cap the two Sam50 barrels on the cytosolic face, forming the SAM^dimer^ complex.^15,16^ From the two Sam50 subunits at the SAM^dimer^ complex, one is a stable subunit that directly interacts with β-barrel precursors while the second shows a dynamic association and acts as a placeholder for incoming substrates. Sam35 sits on top of the stable Sam50 subunit and is essential for mediating the association of the precursor β-signal with the first N-terminal β-strand of Sam50 at the lateral gate, a required step to initiate precursor β-strand assembly at SAM. Sam37 interacts with Sam35 and caps the dynamic Sam50 subunit at the SAM^dimer^ complex. The growing barrel of the precursor displaces the dynamic Sam50 subunit generating a transient SAM^substrate^ complex such that the precursor barrel is now capped by the peripheral subunit Sam37.^15,17,18^ Sam37 contains an a-helical protrusion which, together with the N-terminal segment of the precursor preceding the β-domain, facilitates barrel formation of the precursor.^15,17,18^

An additional form of the sorting and assembly machinery is the SAM^Mdm10^ complex.^15^ In the SAM^Mdm10^ complex, the dynamic Sam50 subunit is replaced by the mitochondrial distribution and morphology protein Mdm10, another β-barrel protein required for the biogenesis of Tom40. During Tom40 biogenesis the a-helical TOM subunits Tom5 and Tom6 assemble with the Tom40 β-barrel while it is still connected to the SAM machinery.^18^ Mdm10 is required for the assembly of the Tom40 module with Tom22 to form the mature TOM core complex. Subsequently Mdm10 itself associates with Sam37 at the SAM machinery to form the SAM^Mdm10^ complex.^11,15,19^ This exchange of β-barrel subunits and β-barrel precursor (SAM^dimer^, SAM^substrate^, SAM^Mdm10^) is termed the β-barrel switching mechanism.^15^ Sam50 contains one N-terminal polypeptide-transport-associated domain (POTRA) which adopts a βααββ fold in the intermembrane space and supports efficient β-barrel protein assembly. ^14,16,20–22^

The β-barrel outer membrane protein Mdm10 is a moonlighting protein. In addition to its role as subunit of the SAM complex, Mdm10 is also a subunit of the ER-mitochondria encounter structure (ERMES), which forms an ER-mitochondria membrane contact site.^23,24^ The ERMES complex consists of the α-helical ER membrane-anchored protein Mmm1, the cytosolic subunits Mdm12 and Mdm34, the β-barrel protein Mdm10 as mitochondrial membrane anchor, and the a-helical outer mitochondrial membrane proteins Gem1(Miro-1) and Tom7.^25–27^ The three subunits Mmm1, Mdm12 and Mdm34 contain synaptotagmin-like mitochondrial-lipid binding protein (SMP) domains, which form extended hydrophobic tunnels for the exchange of lipids between the ER and the mitochondrial outer membrane with preference for phospholipids.^28–30^ Even though Mdm10 is present in SAM and ERMES, both complexes are clearly separated. Isolations of Mmm1 and Mdm12 do not contain other SAM subunits and vice versa, isolations of Sam35 and Sam50 do not contain other ERMES subunits.^31,32^ Moreover, the small TOM protein subunit Tom7 does not bind to SAM^Mdm10^ but only to free Mdm10, thereby regulating the distribution of Mdm10 between the SAM and ERMES complexes.^27,33^ When *TOM7* is deleted the assembly of the TOM complex is enhanced due to the increase of the Mdm10 SAM/ERMES ratio.^27,33,34^

In addition, SAM physically interacts with the mitochondrial import complex (MIM) composed of the two a-helical outer membrane proteins Mim1 and Mim2. The MIM complex is crucial for the biogenesis of a-helical mitochondrial outer membrane proteins including a-helical TOM subunits.^22,34–39^

The detailed functional and structural characterization of the SAM complex suggested that all proteins operating at SAM are known. Here we report, however, that a small α-helical mitochondrial outer membrane protein, initially termed ‘mitochondrial class one protein of 6 kDa’ (Mco6) in our proteomic study of *Saccharomyces cerevisiae* mitochondria,^5^ assembles with the SAM^Mdm10^ complex and cooperates with Mdm10 in the biogenesis of the TOM complex. Based on the identification of this role, we propose to use the functional name ‘Mdm10 complex protein of 6 kDa’ for Mco6.

## RESULTS

### Mco6 forms distinct mitochondrial high molecular mass complexes

Our recent high-resolution mitochondrial blue native PAGE complexome profiling uncovers a co-migration of the *S. cerevisiae* yeast protein Mco6 (Yjl127c-b/YJ127_YEAST) at ~180 kDa with all four known subunits of the sorting and assembly machinery (SAM), Sam35, Sam37, Sam50 and Mdm10 (Figures 1A and S1A).^40^. In addition, Mco6 migrates in a lower molecular mass form of ~100 kDa (Figures 1A and S1A). In our mitochondrial proteome study, Mco6 was identified and classified as mitochondrial outer membrane protein of unknown function.^5^ Subcellular fractions collected by differential centrifugation enrich mitochondrial Tom70 in the pellet of the 13,000xg fraction, Sec61 of the endoplasmic reticulum SEC complex in the 100,000xg fraction and the cytosolic 3-phosphoglycerate kinase involved in glycolysis and gluconeogenesis in the supernatant of the 100,000xg fraction. TwinStrep-tagged Mco6 (Mco6_TwinStrep_) was enriched in the 13,000xg pellet fraction, which confirms the mitochondrial localization of Mco6 predicted by the high confidence mitochondrial proteome study (Figure S1B).^5^ To analyze the assembly of Mco6 in mitochondria, radiolabeled precursor protein was generated by *in vitro* translation in the presence of [^35^S]methionine. *In organello* import of [^35^S]Mco6 into isolated wild-type (WT) mitochondria was performed and a time dependent accumulation of Mco6 was observed (Figure 1B). The assembly of imported Mco6 was followed by blue native PAGE and Mco6 predominantly assembled into complexes of approximately ~300 kDa and ~160 kDa as assessed by the soluble blue native PAGE marker proteins (Figure 1C), which reflects the migration profile of endogenous Mco6 in the complexome profiling data set.^40^

**Figure 1.**
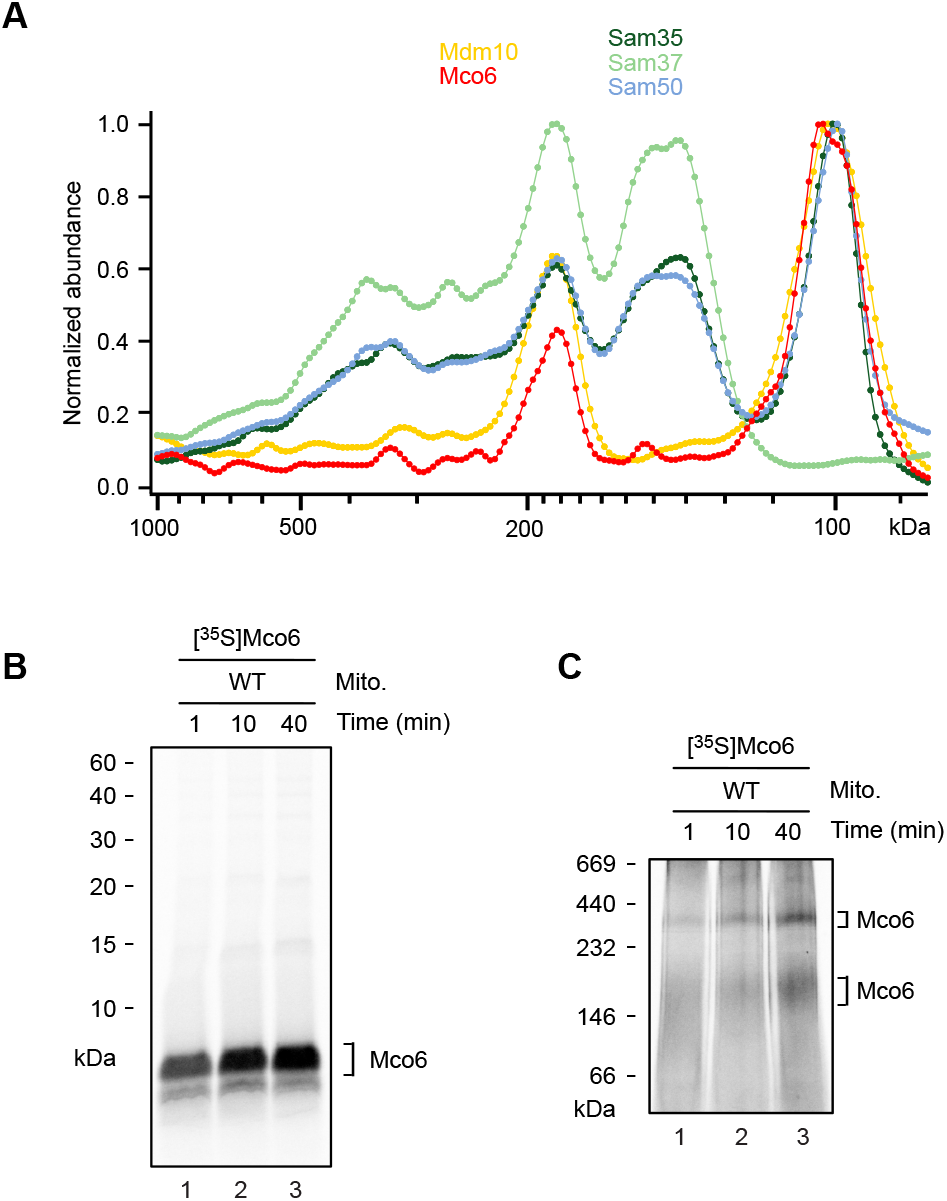
Mco6 assembles into a mitochondrial high molecular mass complex. (A) Mitochondrial blue native PAGE complexome profiles of Mdm10, Mco6, Sam35, Sam37 and Sam50, derived from the complexome data of Schulte et al.^40^ See also Figure S1A. (B,C) Radiolabeled [^35^S]Mco6 was incubated with isolated wild type (WT) mitochondria. At indicated timepoints mitochondria were reisolated, resuspended in SDS sample buffer and analyzed using Tris/Tricine PAGE (B) or solubilized with Digitonin and analyzed using blue native PAGE (C) and subsequent analysis by autoradiography.

### Mco6 assembles into a complex with the sorting and assembly machinery

To analyze if Mco6 is a *bona fide* SAM complex subunit, we compared its blue native PAGE migration in WT mitochondria and Sam50_ΔPOTRA_ mitochondria, where the ~13 kDa polypeptide transport associated (POTRA) domain is deleted.^14,20–22^ The specific size shift of the Mco6-containing complex in Sam50_ΔPOTRA_ to lower molecular mass indicates that Mco6 specifically assembles with the SAM complex (Figure 2A). To test if the assembly of Mco6 with SAM depends on the SAM core subunit Sam37, Mco6 was imported into WT and *sam37*Δ mitochondria.^13^ Mco6 assembly into the SAM complex was completely blocked in *sam37*Δ mitochondria (Figure 2B). In contrast to the transient intermediates of β-barrel precursor proteins like Tom40 and the voltage dependent anion channel Por1 (Porin/VDAC), which form lower molecular mass intermediates with the Sam50-Sam35 subcomplex lacking Sam37, Mco6 does not assemble into a lower mass SAM subcomplex lacking Sam37.^13,41–43^ On the one hand this confirms that Mco6 specifically assembles with the SAM machinery and on the other hand suggests that Mco6 does not assemble with the Sam50-Sam35 subcomplex. We compared the assembly of Mco6 in WT mitochondria and *mdm10*Δ mitochondria.^11^ Similar to the deletion of *SAM37*, Mco6 does not assemble into the SAM complex in *mdm10*Δ mitochondria (Figure 2C). Mitochondria lacking *MDM10* harbor a fully assembled SAM^core^ complex consisting of Sam50, Sam35 and Sam37.^11^ This suggests that Mco6 specifically assembles with the SAM^Mdm10^ complex to form SAM^Mco6^. The a-helical protrusion of Sam37 interacts with luminal residues within the associated β-barrel proteins like Mdm10 (SAM^Mdm10^) and Tom40 (SAM^Tom40^) at the site where the β-barrel substrate strands are assembled (SAM^substrate^).^15,17,18^ The assembly of Mco6 was assessed in *sam37*Δ_186-209_ mitochondria lacking the a-helical protrusion.^17^ The reduced assembly of Mco6 into the SAM complex in *sam37*Δ_186-209_ mitochondria lacking the a-helical protrusion of Sam37 further supports the view that Mco6 specifically assembles with Mdm10 in the SAM^Mco6^ complex (Figure 2D).

**Figure 2.**
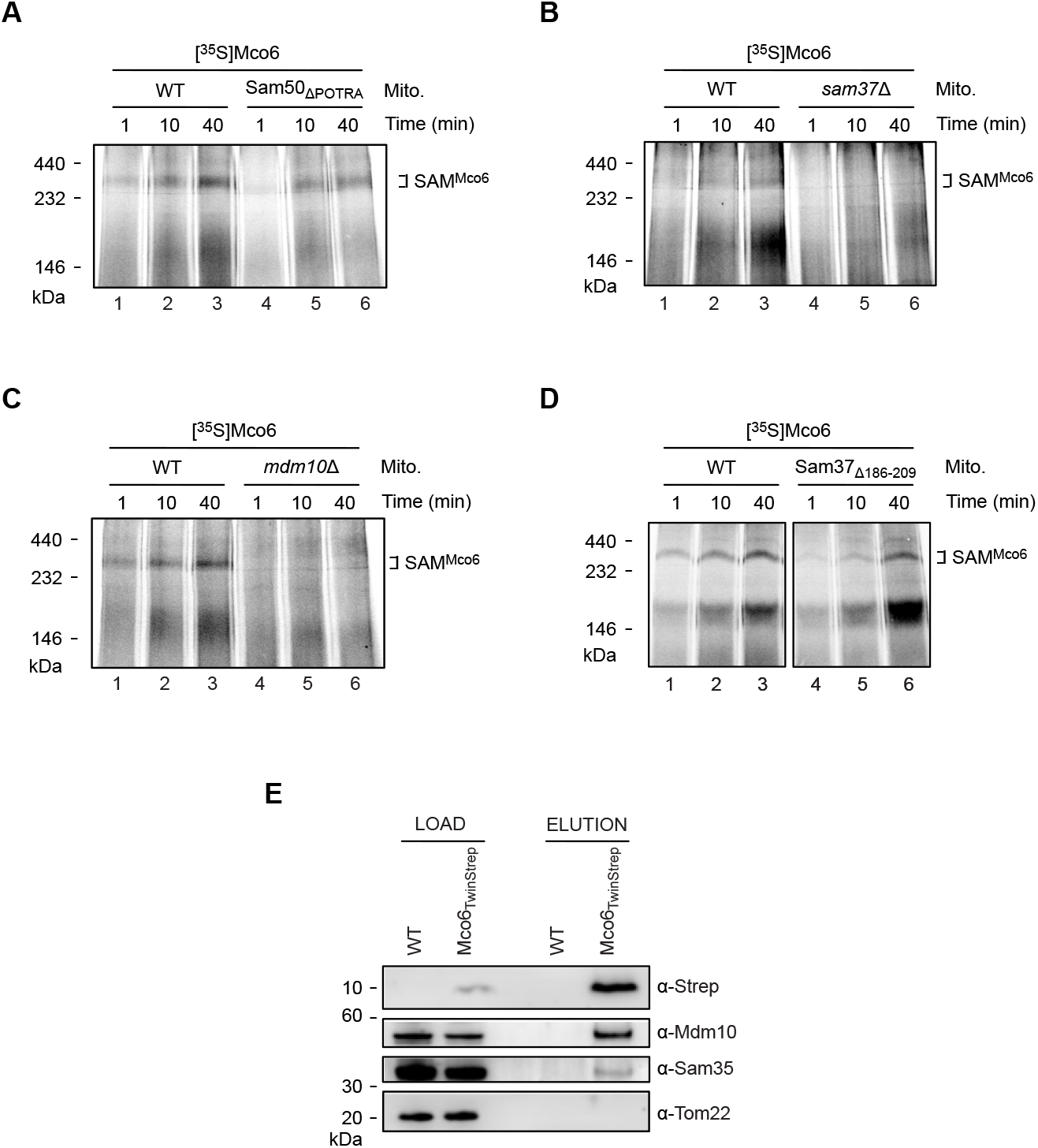
Mco6 assembles together with Mdm10 into the SAM complex. (A-D) Radiolabeled [^35^S]Mco6 was incubated with mitochondria isolated from WT and Sam50_ΔPOTRA_ (Δ2-120) (A), WT and *sam37*Δ (B), WT and *mdm10*Δ (C) or WT and Sam37Δ_186-209_ (D) yeast cells for the indicated timepoints. Reisolated mitochondria were solubilized with digitonin and analyzed by blue native PAGE and autoradiography. (E) Mitochondria isolated from WT and Mco6_TwinStrep_ yeast strains were solubilized with digitonin. Eluates from Strep-tag affinity purifications using magnetic beads (MagStrep type3 XT) were analyzed with NuPAGE gels followed by immunoblotting. Load, 4%; Elution, 100%.

Furthermore, we separated mitochondrial protein complexes by blue native electrophoresis and analyzed single subunits of the complexes by SDS-PAGE as second dimension followed by immunoblotting. The second dimension approach reveals a comigration of the SAM forms of Mdm10 and Mco6_Twinstrep_, supporting the view that Mco6 assembles with the high molecular mass form of the SAM complex with Mdm10 (Figure S2A). To unambiguously demonstrate that Mco6 interacts with the SAM^Mdm10^ complex, we analyzed the interaction partners of Mco6 by affinity purification. Mco6_TwinStrep_ pulldown eluates confirm an interaction of Mco6 with Mdm10 and the SAM^core^ complex revealed by immunoblotting with antibodies against Mdm10 and Sam35 (Figure 2E). Taken together, these results show a stable interaction of Mco6 with the SAM^Mdm10^ complex and, specifically, with its Mdm10 subunit. Therefore, we now refer to the Mco6 & Mdm10-containing SAM complex as SAM^Mco6^.

### Mco6 interacts with, but does not depend on the MIM machinery

An enhanced assembly of Mco6 into the SAM complex was observed in *mco6*Δ mitochondria (Figures 3A and S2B-D). This indicates that endogenous mitochondrial Mco6 and imported radiolabeled [^35^S]Mco6 compete with each other for the binding site on Mdm10. In *mco6*Δ mitochondria the Mco6 complex migrating at ~160 kDa compared to the soluble markers was increased in its intensity, reflecting that this is a specific Mco6-containing complex. We wondered if this complex represents an assembly with the MIM machinery, which is crucial for the membrane insertion of α-helical outer membrane proteins.^36–39,44^ To check if the lower mass Mco6 complex represents an intermediate of Mco6 with the MIM machinery, the assembly of Mco6 was analyzed in WT and _protA_Mim1 mitochondria.^44^ The shift of the Mco6 band in _ProtA_Mim1 mitochondria due to the Protein-A tag on Mim1 confirmed that Mco6 forms in addition to SAM also a complex with the MIM machinery (Figure 3B). A pulse chase import of Mco6 and Tom40 revealed that Mco6 remains stably associated with the SAM and MIM machineries in contrast to Tom40 which is chased into the intermediate II and the mature TOM complex (Figure 3C).^17^ This finding indicates that the association of Mco6 with the SAM^Mdm10^ complex does not represent an assembly intermediate. Mco6 is an interaction partner of the SAM^Mdm10^ complex, which is in exchange with the MIM machinery. To test if the import of Mco6 depends on the MIM machinery itself, the assembly of Mco6 was tested in WT and *mimID* mitochondria.^44^ Even though Mco6 is an a-helical outer membrane protein and thus potentially a bona fide substrate for the MIM machinery, its assembly into the SAM^Mco6^ complex occurs independently of the presence of MIM, while the Mco6-Mim1 complex formation is blocked in its absence (Figure 3D).

**Figure 3.**
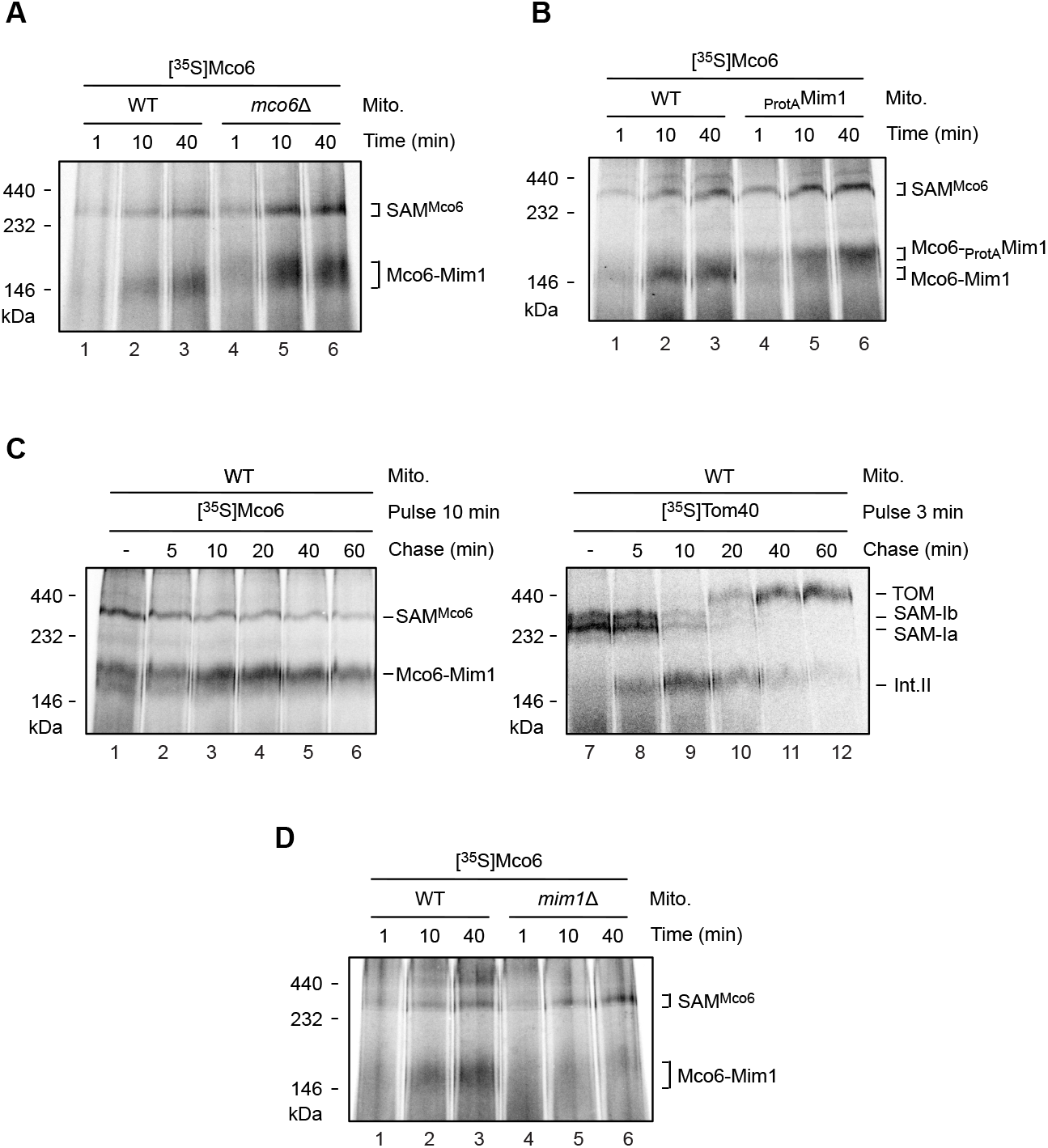
Relation of Mco6 to the MIM machinery. (A,B) Assembly of [^35^S]Mco6 in WT and *mco6*Δ (A) or chromosomal N-terminal ProtA-tagged Mim1 (_protA_Mim1) (B) mitochondria as described for (Figure 2B-E). (C) Radiolabeled [^35^S]Mco6 and [^35^S]Tom40 were incubated with WT mitochondria for 10 and 3 min, respectively (pulse). Mitochondria were reisolated and radiolabeled Mco6 and Tom40 chased for the indicated time periods. Reisolated mitochondria were solubilized with digitonin and analyzed by blue native PAGE and autoradiography. TOM, translocase of the outer mitochondrial membrane; SAM^Mco6^, sorting and assembly machinery containing Mco6; SAM-Ia, sorting and assembly machinery intermediate Ia containing Sam50, Sam35, Sam37 and Tom40; SAM-Ib, sorting and assembly machinery intermediate Ib containing Sam50, Sam35, Sam37, Tom40, Tom5 and Tom6; MIM^Mco6^, mitochondrial import complex containing Mco6; Int.II, intermediate II complex containing Tom40, Tom5, Tom6 and Tom7. (D) Assembly of [^35^S]Mco6 in WT and *mim1*Δ mitochondria by blue native PAGE.

### Mco6 cooperates with Mdm10 in the assembly of the TOM complex

Since Mco6 is directly associated with the SAM^Mdm10^ complex forming the SAM^Mco6^ complex, we tested if Mco6 and Mdm10 display a genetic interaction. Since the deletion of *MDM10* causes several pleiotropic effects like loss of mitochondrial DNA and altered mitochondrial morphology, we selected the *mdm10* yeast mutant *mdm10-27*, which is able to grow on respiratory media, for genetic studies (Figure 4A). Growth analysis of WT, *mco6Δ, mdm10-27* and the *mdm10-27 mco6*Δ double mutant revealed a strong negative genetic interaction between *MCO6* and *MDM10* (Figure 4A). In isolated mitochondria of the *mdm10-27 mco6*Δ double mutant, a significant reduction of Tom22 and of the TOM complex was observed (Figure 4B and C), which was not observed in Mdm10-ts27 mitochondria (Figure S3A and B). Moreover, deletion of *MCO6* causes a temperature-sensitive growth defect at elevated temperature, which is especially pronounced on non-fermentable glycerol medium (YPG) (Figure 4D). To analyze the defects of mitochondria lacking Mco6, precultures of WT and *mco6*Δ yeast strains were cultivated at permissive temperature and shifted to the non-permissive 39°C conditions for 72h. This caused a growth retardation of *mco6*Δ compared to WT. Protein levels of isolated mitochondria from cells shifted to non-permissive conditions were analyzed with specific focus on mitochondrial outer membrane proteins. In contrast to the essential β-barrel membrane proteins Tom40 and Sam50 and the most abundant outer membrane β-barrel proteins Por1, only the levels of Mdm10 were specifically reduced in mitochondria lacking Mco6 (Figure 4E). In addition, protein levels of the mitochondrial import complex (MIM) were also slightly reduced (Figure 4E). To analyze the relation between Mdm10 and the MIM machinery, protein levels of Mdm10 were analyzed in *mim1*Δ mitochondria and protein levels of Mim1 and Mim2 were analyzed in *mdm10*Δ mitochondria, respectively.^11,44^ Levels of Mdm10 comparable to WT were observed in *mim1*Δ mitochondria (Figure S4A). In contrast, slightly reduced levels of Mim1 and Mim2 were detected in *mdm10*Δ mitochondria (Figure S4B). This analysis shows that the stability of Mdm10 is specifically affected by the deletion of *MCO6* at non-permissive conditions and suggests that the reduced levels of the MIM machinery subunits Mim1 and Mim2 might at least partially be a consequence of the decreased Mdm10 levels (Figure 4E). The reduced levels of Mdm10 in *mco6D* at non-permissive conditions causes assembly defects of Tom22 and Tom5 into the TOM complex (Figures 4F and G). In contrast, TOM assembly is comparable to WT in mitochondria isolated from cells cultivated at permissive conditions (Figures S2C, S2D, S4C and S4D). The TOM assembly defects observed in *mco6D* mitochondria are in line with previous results in *mdm10D* mitochondria.^11^ These results demonstrate that Mco6 is specifically associated with Mdm10 forming the SAM^Mco6^ complex (Figure 4H), which is in agreement with Colab Alphafold2 modelling of Mco6 with Mdm10.^45^ We conclude that Mco6 is required for the assembly of the TOM complex.

**Figure 4.**
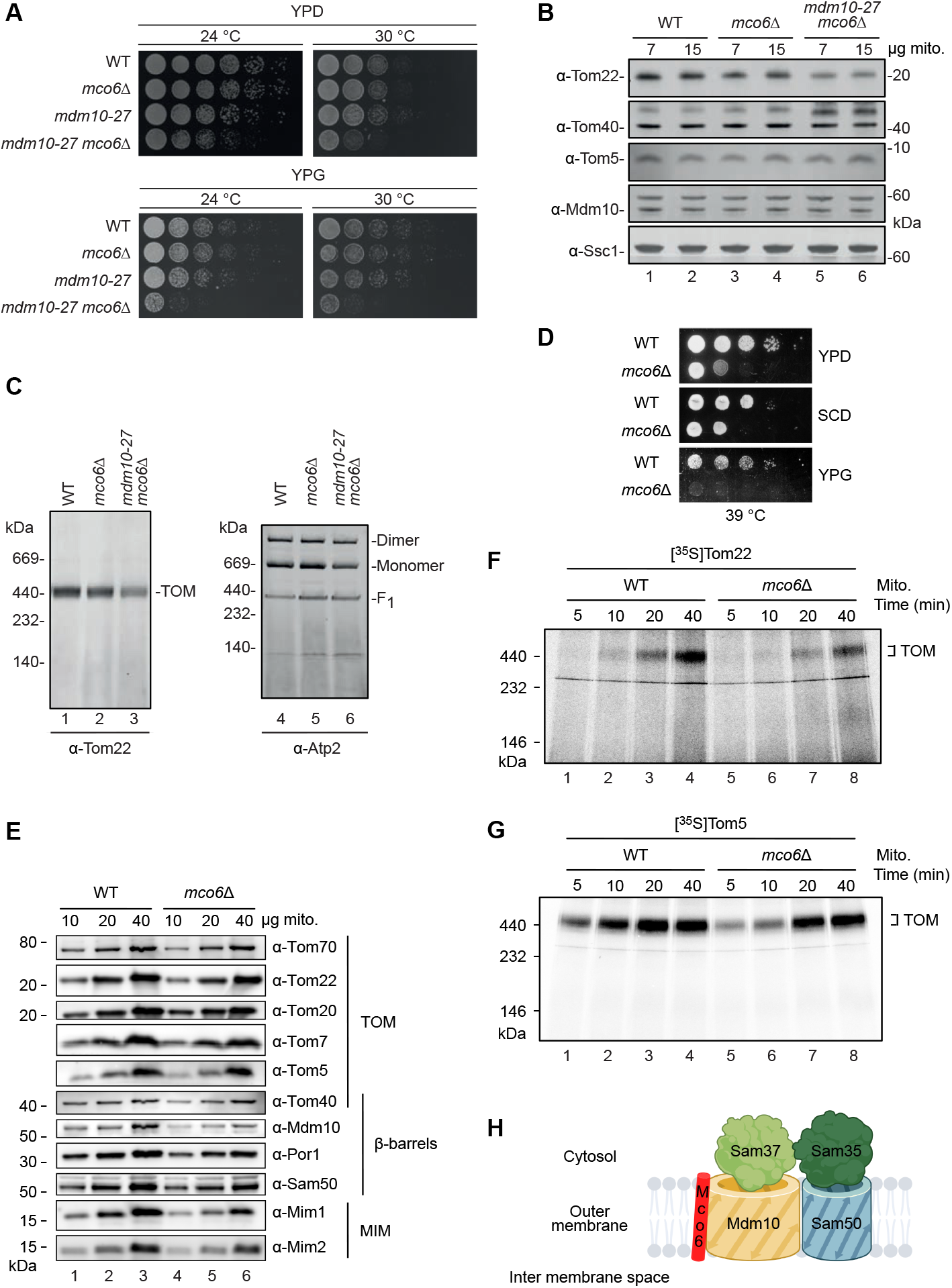
Mco6 is required for maintenance of Mdm10. (A) Growth assay with serial dilutions of wild type (WT), *mco6*Δ, *mdm10-27* and *mdm10-27 mco6*Δ yeast cells on agar plates containing fermentable glucose (D, dextrose) and non-fermentable glycerol (G) medium at 24°C and 30°C. YPD, yeast extract-peptone-dextrose medium; YPG, yeast extract-peptone-glycerol medium. (B) Protein levels of mitochondria isolated from WT, *mco6*Δ, and *mdm10-27 mco6*Δ yeast cells. The indicated total mitochondrial protein amounts were resuspended in SDS sample buffer and analyzed using MES-SDS PAGE and immunoblotting. (C) Protein complexes of mitochondria isolated from WT, *mco6*Δ, and *mdm10-27 mco6*Δ yeast cells. Samples were solubilized with digitonin and analyzed by blue native PAGE and immunoblotting. TOM, translocase of the outer mitochondrial membrane; F1, Monomer and Dimer, Different assembly states of Atp2 into the F1 intermediate or the different forms of the mitochondrial F_O_F_1_-ATPase (Complex V). (D) Growth assay with serial dilutions of WT and *mco6*Δ yeast cells on agar plates containing fermentable glucose (D, dextrose) and non-fermentable glycerol (G) medium at 39°C. SCD, synthetic complete-dextrose medium. (E) Protein levels of mitochondria isolated from WT and *mco6*Δ yeast cells cultured with YPG medium after shifting cells to non-permissive conditions (39 °C for 72 hours). The indicated total mitochondrial protein amounts were resuspended in SDS sample buffer and analyzed using Tris/Tricine PAGE and immunoblotting. (F,G) Radiolabeled [^35^S]Tom22 (F) or [^35^S]Tom5 (G) was incubated with mitochondria isolated from WT and *mco6*Δ yeast cells grown in YPG medium at non-permissive conditions (39 °C for 72 hours) for the indicated timepoints. Samples were then solubilized with digitonin and analyzed using Blue Native PAGE and autoradiography. (H) Model of the SAM^Mco6^ complex (created with BioRender.com based on Mco6 Colab Alphafold2 modelling with Mdm10 and PDB 7BTX).^15^

## DISCUSSION

Mitochondria evolved by endosymbiosis from a Gram-negative bacterial ancestor and therefore the mitochondrial outer membrane contains β-barrel membrane proteins. These β-barrel membrane proteins are assembled into large oligomeric complexes. The β-barrel Tom40 assembles with the a-helical membrane proteins Tom5, Tom6, Tom7, Tom22 and interacts with the receptors Tom20, Tom70 and Om14.^40,46–48^ After identification of the a-helical membrane proteins of the TOM complex, the specific complex formation between β-barrel membrane proteins and a-helical membrane proteins was regarded as a special case. Meanwhile the porin β-barrel membrane protein was found to interact with the a-helical membrane proteins Om14 and Om45 and the β-barrel protein Mdm10 interacts as subunit of the ER-mitochondria encounter structure (ERMES) with the a-helical outer membrane proteins Tom7 and Gem1(Miro-1)^23,25–27,49^

Here we demonstrate that the a-helical outer membrane protein Mco6 assembles with the Mdm10 form of the sorting and assembly machinery to form the ~180 kDa SAM^Mco6^ complex composed of Sam50, Sam37, Sam35, Mdm10 and Mco6 (54,4 kDa; 37,5 kDa; 37,4 kDa; 56,2 kDa and 6,0 kDa, respectively) (Figures 1A, 2, 4H and S2A). Thus, in contrast to bacteria, mitochondrial β-barrel membrane proteins have numerous α-helical membrane proteins as specific interaction partners. We report by two independent lines of evidence that Mco6, in cooperation with Mdm10, is required for the assembly of the TOM complex. On the one hand, *MCO6* has a negative genetic interaction with the temperature-sensitive mutant *mdm10-27* (Figure 4A). Isolated double mutant mitochondria have reduced levels of Tom22 and assembled TOM complex (Figures 4B and 4C). On the other hand, deletion of *MCO6* causes a temperature-sensitive growth defect and, in isolated mitochondria, the protein levels of the Mco6 partner protein Mdm10 are specifically affected, which causes assembly defects of the TOM complex (Figures 4D–4G). Even before Mco6 was identified as high confidence mitochondrial outer membrane protein, Costanzo et al. reported a negative genetic interaction between *MCO6* and *TOM7*.^5,50^ Our results show that Mdm10 not only specifically interacts with Tom7 but also with Mco6. The negative genetic interaction supports the view that both smORF proteins (encoded by small open reading frames) Tom7 and Mco6 control the function of Mdm10.

Rasul et al. reported that the Mco6 homolog Emr1 of *S. pombe* regulates the number of Mdm12 ERMES foci and concluded that Emr1 interacts with Mdm12.^51^ Our finding that Mco6 specifically interacts with the Mdm10 form of the sorting and assembly machinery (SAM^Mdm10^), which does not contain any other ERMES core subunits Mmm1, Mdm12 and Mdm34, suggests that Emr1/Mco6 also likely interacts specifically with Mdm10 as subunit of the ERMES complex. This provides a reasonable explanation for the reduction of ERMES foci, the mitochondrial morphology defects and the impaired mitochondrial segregation observed by Rasul et al.^51^

Moreover, we provide evidence that Mco6 also associates with Mim1 of the MIM complex, which is itself required for the biogenesis of a-helical outer membrane proteins (Figure 3).^36,39^ The specific sharing of interaction partners between different outer membrane complexes may provide a mechanism to regulate the functional network of machineries that coordinate mitochondrial metabolism, protein and lipid biogenesis. It is tempting to speculate that our study about Mco6 uncovered such a novel regulatory protein.

Taken together, our results suggest that Mco6 specifically cooperates with Mdm10 in the mitochondrial sorting and assembly machinery, forming the SAM^Mco6^ complex. This cooperation is required for the efficient assembly of the main mitochondrial protein entry gate TOM.

## METHODS

### Yeast strains, plasmids and primers

The *Saccharomyces cerevisiae* yeast strains, plasmids and primers are provided in Tables S1–S3. Deletion of *MCO6* was performed using homologous recombination with a kanamycin resistance cassette as described.^52^ Chromosomal C-terminal TwinStrepII tagging of Mco6 was performed using homologous recombination following the same procedure as for *MCO6* deletion. Correct integration of the TwinStrepII tag was verified by PCR and sequencing. Used plasmid templates and primers are listed in Supplementary Tables S2 and S3.

### Yeast growth assays

The growth phenotype of yeast strains was assessed using the drop-test approach on agar plates containing yeast extract-peptone-dextrose medium (YPD), synthetic complete-dextrose medium (SCD) or yeast extract-peptone-glycerol medium (YPG) at the indicated growth temperatures. Yeast strains were grown overnight in YPD medium at 30°C, diluted to an optical density (OD_600_) of 0.3 with fresh YPD and grown further at 30°C for 3-5 hours. One ml of an OD_600_ = 1.0 from each strain were washed twice with H_2_O to completely remove the growth medium before plating. Ten-fold serial dilutions were finally performed starting with an OD_600_ = 1.0 and 3 μl per drop were plated using a multichannel pipette. Cells were grown at the respective temperatures for 1 to 8 days.

### Mitochondria isolation

Yeast cells were grown in yeast extract-peptone-glycerol medium (YPG) (2% (w/v) bacto-peptone, 1% (w/v) yeast extract and 3% glycerol) or YPS medium (with 2% (w/v) sucrose) at 30°C or 39°C with shaking at 130 rpm. At early logarithmic phase, mitochondria were isolated by differential centrifugation as described previously.^53^ Briefly, yeast cells were harvested by centrifugation, washed in water and subsequently incubated in prewarmed DTT softening buffer (100 mM Tris/H_2_SO_4_, pH 9.4, 10 mM DTT) at the respective growth temperature. Yeast cell wall was disrupted by incubation with zymolyase buffer (16 mM K_2_HPO_4_, 3.96 mM KH_2_PO_4_, pH 7.4, 1.2 M sorbitol (Roth)) in the presence of 5 mg/g zymolyase at the respective growth temperature. Spheroblasts were then washed with zymolyase buffer (without zymolyase) and opened mechanically on ice with a homogeniser in cold homogenisation buffer (0.6 M sorbitol, 10 mM Tris/HCl, pH 7.4, 1 mM ethylenediaminetetraacetic acid (EDTA), 0.4% [w/v] bovine serum albumin (Sigma), 1 mM phenylmethyl sulfonyl fluoride (PMSF, Sigma)). Cell debris were removed by centrifugation and mitochondria finally isolated by centrifugation at 16,000xg, 15 min, 4°C. Mitochondrial pellets were washed in ice cold SEM buffer (10 mM MOPS/KOH, pH 7.2, 250 mM sucrose, 1 mM EDTA), reisolated and resuspended in SEM buffer to a protein concentration of 10 mg/ml. Mitochondria were finally snap-frozen as aliquots in liquid nitrogen and stored at −80°C. To isolate mitochondria from the *mco6*Δ yeast strain at non-permissive temperature, the cells were pre-grown at permissive temperature (30°C) in YPG medium, diluted 60x into pre-warmed YPG medium at 39°C and further grown for 72h at 39°C before mitochondria isolation as described above.

### Subcellular fractionation

Subcellular fractionation was performed using 70 OD of yeast cells grown to a OD_600_ of 1. Cells were harvested and washed in dH_2_O. The cell pellet was resuspended in 2 ml DTT buffer (100 mM Tris-H_2_SO_4_ pH 9.4, 10 mM DTT) and incubated for 20 min at 30°C (130 rpm). Subsequently, spheroplasts were generated by incubation in 1 ml zymolyase buffer (1.2 M sorbitol, 20 mM potassium phosphate, pH 7.4, 0.5% [w/v] zymolyase) for 45 min (130 rpm) at 30°C. Spheroplasts were harvested by centrifugation (3,000xg, 5 min). To remove zymolyase a washing step in 1 ml 1.2 M sorbitol was performed. The following steps were performed at 4°C. Spheroplasts were resuspended in 2 ml homogenization buffer (0.6 M sorbitol, 10 mM Tris-HCl, 1 mM EDTA, 1 mM PMSF, pH 7.4) and dounced 20 times in a cooled glass-teflon potter on ice. The samples were subjected to a clearing spin (300xg, 4°C, 5 min, S300). Half of the S300 supernatant was saved as PNS (post nuclear supernatant). The other half of the S300 fraction was subjected to further centrifugation (13,000xg, 15 min, 4°C) to generate the pellet P13. The supernatant fraction (S13) was centrifuged (100,000xg, 60 min, 4°C) to collect the pellet P100 and the supernatant S100. Proteins in the PNS and S100 fractions were precipitated by addition of 25% (v/v) of 50% TCA (final concentration 10%) and incubation for 30 min on ice. Precipitated proteins were pelleted by centrifugation (18,200xg 15 min, 4°C) and the protein pellet was washed once with ice cold Tris base (1M Tris). All samples with 2x Laemmli buffer (120 mM Tris-HCl, pH 6.8, 4% (w/v) SDS, 20% glycerol, 40 mM DTT, 0.1% (w/v) bromphenol-blue) were incubated at 60°C (1,400 rpm, 20 min) and analyzed by SDS PAGE followed by immunoblotting.

### Gel electrophoresis

To analyze mitochondrial protein levels isolated mitochondria were sedimented and resuspended in 2x Laemmli buffer containing 2% beta-mercaptoethanol and 2 mM PMSF. The indicated total mitochondrial protein amounts were loaded into 10% Tris/Tricine gels and the specific protein levels analyzed by immunoblotting.

For blue native polyacrylamide gel electrophoresis (blue native PAGE) isolated yeast mitochondria (50-100 μg) in SEM buffer were pelleted, resuspended in 50 μl of 1% (w/v) digitonin-containing lysis buffer (0.1 mM EDTA, 10% [v/v] glycerol, 50 mM NaCl, 2 mM PMSF, 20 mM Tris/HCl, pH 7.4) and incubated for 15 min on ice. After solubilization, 5.5 μl of a 10x blue native loading dye solution (final concentration: 50 mM ε-Aminocaproic acid, 10 mM bis-tris, 0.5% Coomassie G, pH 7.0) were added to the solubilized mitochondria and insoluble material was removed by centrifugation (10 min, 20,800 × *g*, 4°C). Solubilized mitochondria were finally loaded into gradient 6-16.5% blue-native PAGE gels and run at 600V, 20 mA and 90 min.^54^ The gels were later analysed either by autoradiography or immunoblotting. For two-dimensional gel electrophoresis mitochondria isolated from the Mco6_TwinStrep_ strain (Table S1) were solubilized in 1% (w/v) digitonin-containing lysis buffer as described above and mitochondria protein complexes first separated by using a gradient 6-16.5% blue native PAGE (1 ^st^ dimension). The blue native gel lane was excised and horizontally mounted into a 10% Tris/Tricine gel (2^nd^ dimension). The indicated mitochondrial proteins and protein complexes were finally analysed by immunoblotting. Isolated mitochondria from the Mco6_TwinStrep_ strain were sedimented, resuspended in 2x Laemmli buffer containing 2% beta-mercaptoethanol and 2 mM PMSF and loaded as control in the 2^nd^ dimension (Mito.).

### Affinity purification using the TwinStrepII tag

Affinity purification was performed with MagStrep type3 XT beads (IBA Lifesciences) as recommended by the manufacturer. Briefly, 900 μg isolated mitochondria were harvested, resuspended in solubilization buffer (1x buffer XT, 1% digitonin, 2 mM PMSF, protease inhibitor cocktail and 10% glycerol) and solubilized for 30 min at 4°C with head over head rotation. A small fraction of solubilized mitochondria was then separated and loaded with Laemmli buffer (final 1x Laemmli buffer supplemented with 1% beta-mercaptoethanol and 1 mM PMSF). Solubilized mitochondria were then incubated with pre-washed MagStrep type3 XT beads and incubated for 2h at 4°C with head over head rotation. Unbound material was removed by washing the samples 5x for 5min with head over head rotation at 4°C in wash buffer plus detergent (1x buffer XT, 1% digitonin, 2 mM PMSF and 10% glycerol) using a magnetic separator (New England Biolabs). Bound material was finally eluted with Biotin elution buffer (1x buffer Biotin XT elution, 0,3% digitonin, 2 mM PMSF and 10% glycerol) for 1h at 4°C with head over head rotation. The eluates were finally separated from the beads using the magnetic separator and loaded with Laemmli buffer (final 1x Laemmli buffer supplemented with 1% beta-mercaptoethanol and 1 mM PMSF). Samples were separated using pre-casted Bis-Tris 4-12% NuPAGE gels (Invitrogen) and analyzed by immunoblotting.

### Radiolabeled protein import into isolated mitochondria

^35^S-radiolabeled Tom40 precursor proteins were synthesized *in vitro* using the TNT Quick Coupled Transcription/Translation kit (Promega). Transcripts for *MCO6* encoding 3 additional N-terminal methionine residues for radiolabelling of the Mco6 protein which lacks internal methionine residues, *TOM22* and *TOM5* were generated using the mMESSAGE mMACHINE transcription kit (Thermo Fisher) and finally radiolabelled precursor proteins were synthesized with the Flexi rabbit reticulocyte lysate system (Promega). ^35^S-radiolabeled precursor proteins were incubated with isolated mitochondria in import buffer (3% (w/v) bovine serum albumin, 250 mM sucrose, 80 mM KCl, 5 mM MgCl_2_, 5 mM methionine, 10 mM MOPS/KOH, pH 7.2, 2 mM KH2PO4) containing 4 mM ATP, 4 mM NADH, 5 mM creatine phosphate and 100 μg/ml creatine kinase at 25°C and 300 rpm for the indicated time periods. Import reactions were stopped by transferring the samples to ice. Mitochondria were reisolated, washed with SEM buffer and solubilized with 1% (w/v) digitonin (MATRIX BioScience) in lysis buffer (0.1 mM EDTA, 10% [v/v] glycerol, 50 mM NaCl, 2 mM PMSF, 20 mM Tris/HCl, pH 7.4) for 15 min on ice. Insoluble material was removed by centrifugation (10 min, 20,800×g, 4°C) and protein complexes were analysed by gradient 6-16.5% blue-native PAGE and autoradiography.

### Pulse-chase experiment

^35^S-radiolabeled Mco6 and Tom40 were independently incubated with isolated mitochondria for 10 min and 3 min in import buffer (pulse). Mitochondria were then reisolated, washed with SEM buffer, resuspended in pre-warmed chase buffer (250 mM sucrose, 6 mM EDTA, 4 mM ATP, 10 mM MOPS pH 7.2) and further incubated at 25°C and 300 rpm in a single reaction tube (chase). Aliquots were taken at the indicated time points and samples were directly taken to the next steps. Mitochondria were reisolated, washed, solubilized with 1% digitonin and analysed by blue-native PAGE and autoradiography as described previously.

## SUPPLEMENTAL INFORMATION

Supplemental information Figures S1 to S4 and Tables S1 to S3 can be found below.

## ACKNOWLEDGEMENTS

We thank N. Pfanner for discussion. Work included in this study has also been performed in partial fulfilment of the requirements for the master thesis of H.M. (University of Bonn). This work was supported by the Deutsche Forschungsgemeinschaft (DFG, German Research Foundation) SFB 1381 project ID 403222702 (U.S., B.F. and N.W.); FA 332/15-1 project ID 439189341, FA 332/17-1 project ID 446245862 and TRR 152 project ID 239283807 (B.F.); BE 4679/2-2 project ID 269424439 and SFB 1218 project ID 269925409 (T.B.); WI 4506/1-1 project ID 406757425 (N.W.); Germany’s Excellence Strategy (CIBSS EXC-2189 project ID 390939984, U.S., B.F. and N.W.); and the European Research Council (ERC) Consolidator grant no. 648235 (N.W.).

## AUTHOR CONTRIBUTIONS

N.W. and T.B. conceived and supervised the project. J.B.V, H.M., C.S., E.F.S., S.P.S., L.E., J.S., S.B.S., P.L., B.G., F.d.B., U.S., B.F., T.B. and N.W. designed, performed and analyzed experiments. J.B.V., H.M., E.F.S., B.F. and T.B. prepared figures. J.B.V. and N.W. wrote the manuscript with the support of all of the authors.

---

**Figure S1.**
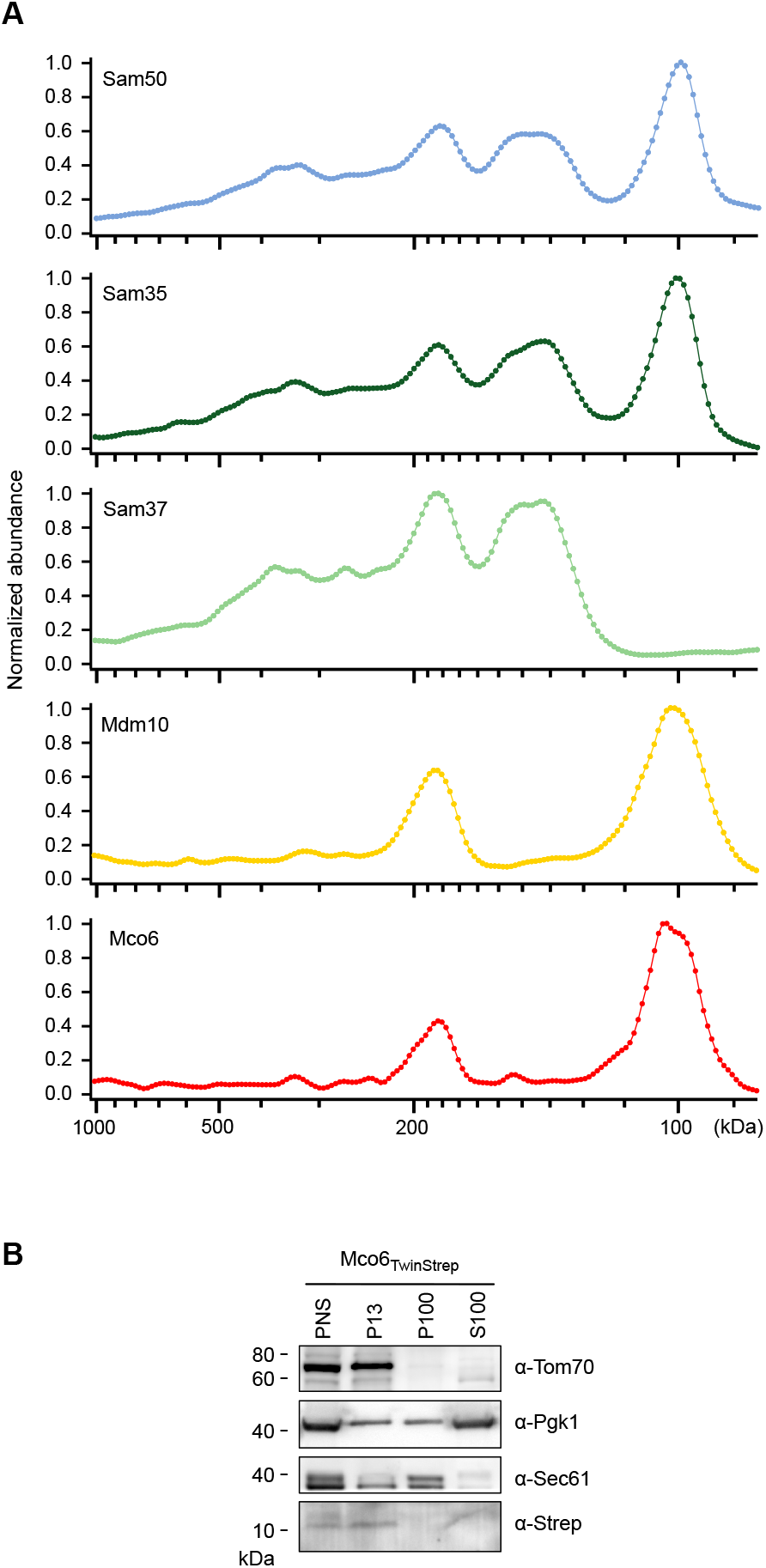
Mco6 complex and subcellular fractionation of Mco6. (A) Individual blue native PAGE yeast complexome profiles of Sam50, Sam35, Sam37, Mdm10 and Mco6 related to Figure 1A, derived from the complexome data of Schulte et al. (Extended Data Figures 5 and 8a and Supplementary Tables 1 and 2, see: http://www.complexomics.org).^40^ (B) Subcellular fractionation of yeast cells with C-terminal TwinStrep-tagged Mco6 (Mco6_TwinStrep_) grown in medium containing non-fermentable glycerol (YPG) at 30°C. The different fractions were resuspended in SDS sample buffer and analyzed using Tris/Tricine PAGE and immunoblotting. PNS, post-nuclear supernatant; P13, mitochondrial fraction; P100, microsomal fraction, S100, cytosolic fraction.

**Figure S2.**
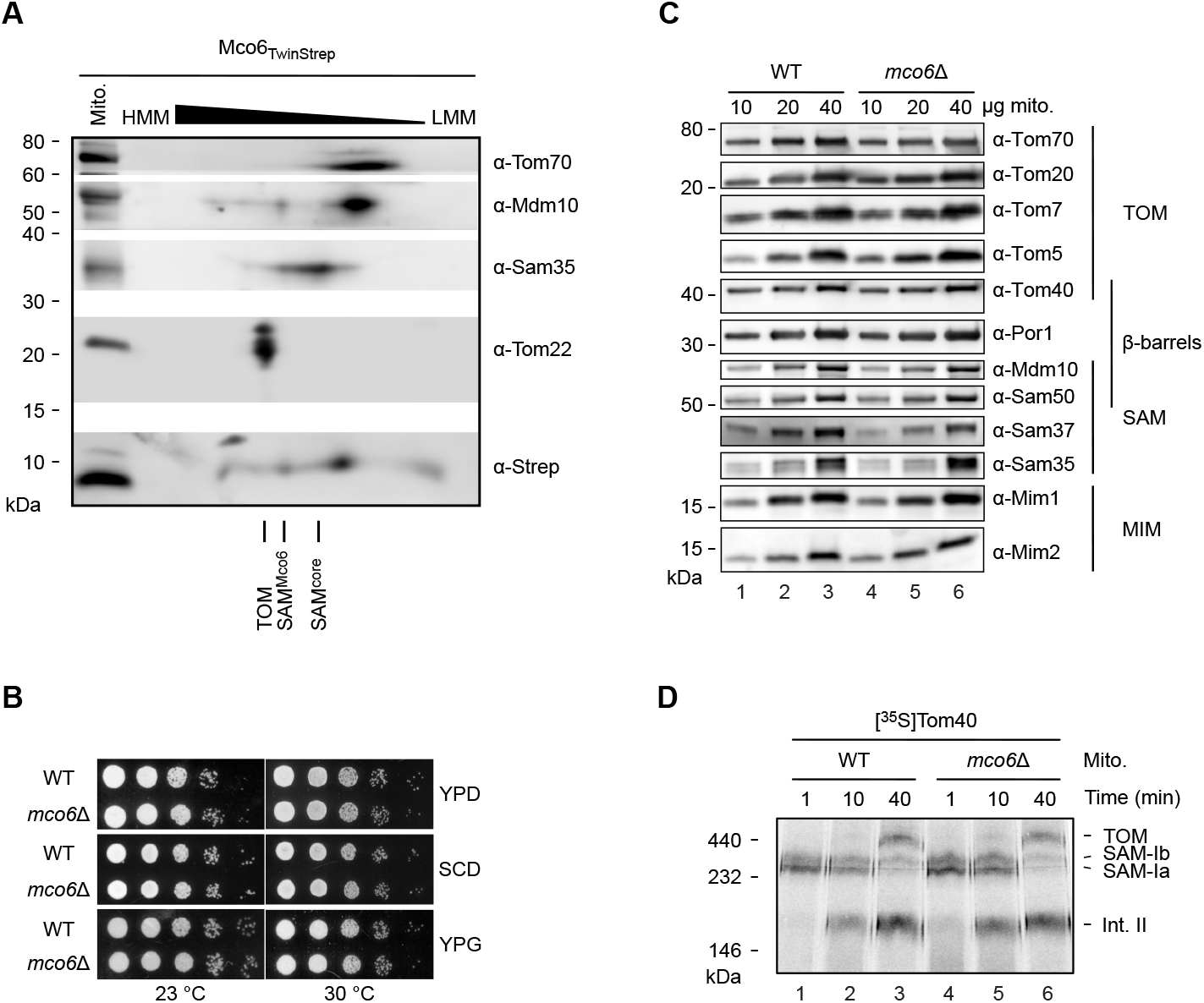
Mco6 complex and analysis of *mco6*Δ mitochondria. (A) Two-dimensional gel of mitochondria isolated from C-terminal TwinStrep-tagged Mco6 (Mco6_TwinStrep_) yeast cells. Mitochondria solubilized with digitonin were analyzed by blue native PAGE (1^st^ dimension). The blue native PAGE lane was then excised, subjected to Tris/Tricine PAGE (2^nd^ dimension) and analyzed via immunoblotting. Additionally, Mco6_TwinStrep_ mitochondria were resuspended in SDS sample buffer and loaded as control for the 2^nd^ dimension (Mito.). HMM, high molecular mass respective LMM, low molecular mass from the 1^st^ dimension. TOM, translocase of the outer mitochondrial membrane; SAM^Mco6^, sorting and assembly machinery containing Mco6; SAM^core^, sorting and assembly machinery containing Sam50, Sam35 and Sam37. (B) Growth assay of serial dilutions of WT and *mco6*Δ yeast cells on agar plates containing fermentable glucose (D, dextrose) and non-fermentable glycerol (G) medium at 23°C and 30°C. YPG, yeast extract-peptone-dextrose medium; SCD, synthetic complete-dextrose medium; YPG, yeast extract-peptone-glycerol medium. (C) Protein levels of mitochondria isolated from WT and *mco6*Δ yeast cells cultured with YPG medium at 30°C. The indicated total mitochondrial protein amounts were resuspended in SDS sample buffer and analyzed using Tris/Tricine PAGE and immunoblotting. (D) Radiolabeled [^35^S]Tom40 was incubated with mitochondria isolated from WT and *mco6*Δ yeast cells for the indicated timepoints. Reisolated mitochondria were solubilized with digitonin and analyzed using blue native PAGE and autoradiography. TOM, translocase of the outer mitochondrial membrane; SAM-Ia, sorting and assembly machinery intermediate Ia containing Sam50, Sam35, Sam37 and Tom40; SAM-Ib, intermediate Ib containing Sam50, Sam35, Sam37, Tom40, Tom5 and Tom6; Int.II, intermediate II complex containing Tom40, Tom5, Tom6 and Tom7.

**Figure S3.**
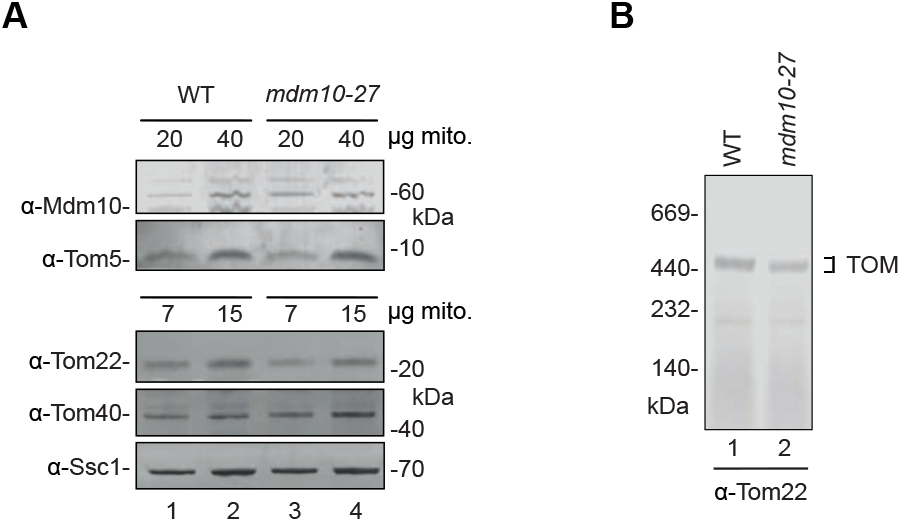
Analysis of *mdm10-27* mitochondria. (A) Protein levels of mitochondria isolated from WT and *mdm10-27* yeast cells. The indicated total mitochondrial protein amounts were resuspended in SDS sample buffer and analyzed using MES-SDS PAGE and immunoblotting. (B) Protein complexes of mitochondria isolated from WT and *mdm10-27* yeast cells. Samples were solubilized with digitonin and analyzed by blue native PAGE and immunoblotting.

**Figure S4.**
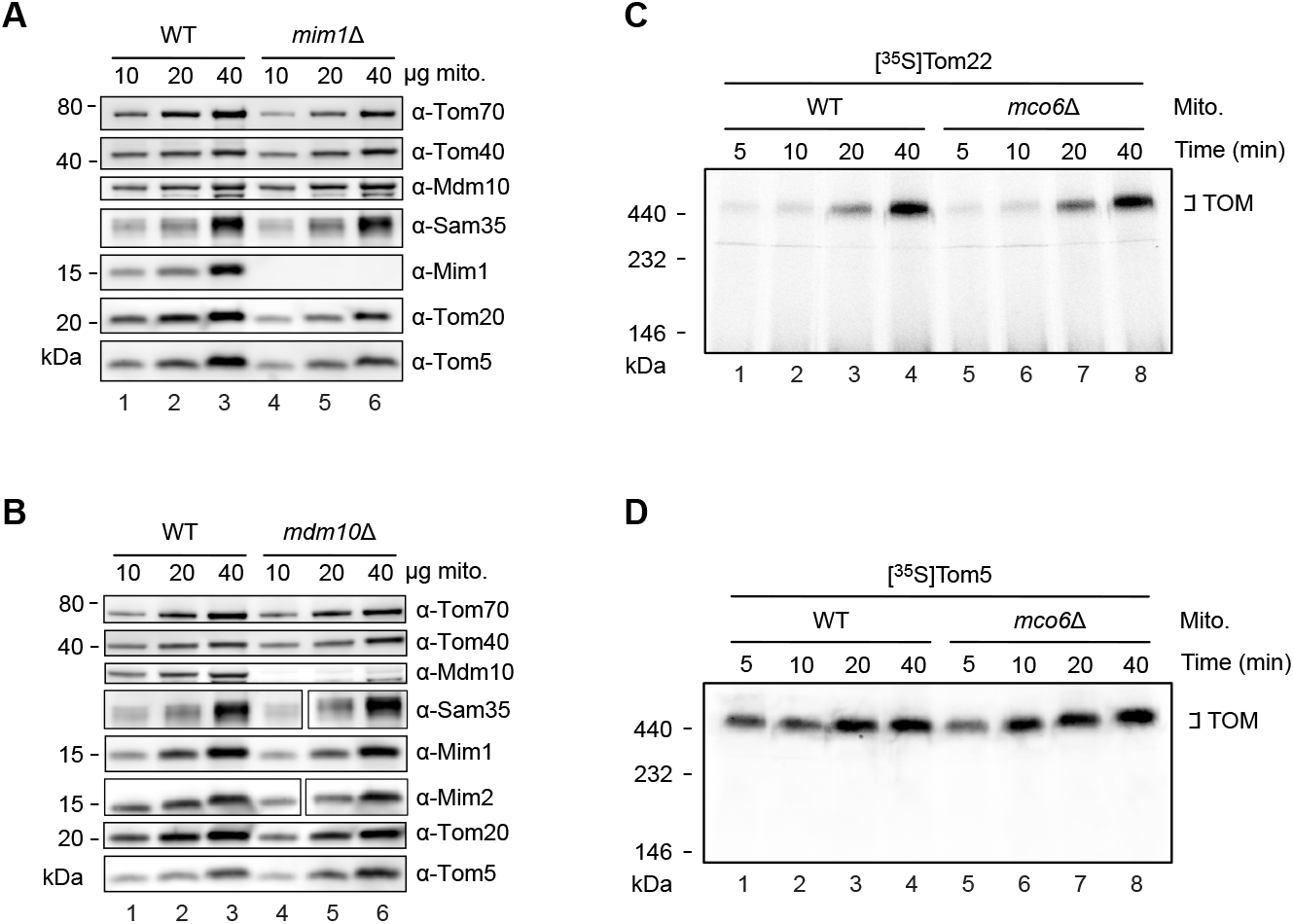
Comparison of WT, *mco6D, mimID* and *mdm1*Δ mitochondria. (A,B) Protein levels of mitochondria isolated from WT and *mim1*Δ (A) or WT and *mdm10*Δ (B) yeast cells. The indicated total mitochondrial protein amounts were resuspended in SDS sample buffer and analyzed using Tris/Tricine PAGE and immunoblotting. (C,D) Radiolabeled [^35^S]Tom22 (C) or [^35^S]Tom5 (D) was incubated with mitochondria isolated from wild type (WT) and *mco6*Δ yeast cells grown in YPG medium at permissive temperature (30°C) for the indicated timepoints. Samples were then solubilized with digitonin and analyzed by blue native PAGE and autoradiography.

**Table S1:** List of *S. cerevisiae* yeast strains used in this study.

| Strain | Reference number | Genotype | Source |
| --- | --- | --- | --- |
| YPH499 wild type (WT) | 1501 | *MATa ura3-52 lys2-801_amber ade2-101_ochre trp1-Δ63 his3-Δ200 leu2-Δ1* |  |
| BY4741 wild type (WT) | 1354 | *MATa his3Δ1 leu2Δ0 met15Δ0 ura3Δ0* | Euroscarf |
| Mco6<sup>TwinStrep</sup> |  | YPH499 *Mco6-TwinStrep::kanMX4* | This study |
| *mco6Δ* | 5699 | BY4741 *mco6::kanMX4* | This study |
| *sam37Δ* | 5647 | YPH499 *sam37::natNT2* | Takeda et al.<sup>17</sup> |
| *mdm10Δ* | 1506 | BY4741 *mdm10::kanMX4* | Euroscarf |
| *mim1Δ* | 2195 | YPH499 *mim1::ADE2* | Becker et al.<sup>44</sup> |
| Sam50<sup>ΔPOTRA</sup> | 2631 | YPH499 *sam50::ADE2 [pFL39-pSam50-Sam50Δ2-120-Sam50t (TRP1)]* | Kutik et al.<sup>14</sup> |
| <sup>ProtA</sup>Mim1 | 1938 | BY4741 *mim1::HIS3-pNOP1-ProtA-TEV-MIM1* | Becker et al.<sup>44</sup> |
| Sam37Δ186-209 | 5648 | YPH499 *Sam37Δ186-209(GAG linker)::hphNT1* | Takeda et al.<sup>17</sup> |
| WT (for Mdm10 strains) | 3402 | YPH499 *mdm10::ADE2 pFL39-MDM10WT(TRP1)* | Flinner et al.<sup>55</sup> |
| Mdm10 shuffle |  | YPH499 *mdm10::ADE2 pYep352-MDM10(URA3)* | Flinner et al.<sup>55</sup> |
| *mco6Δ* |  | YPH499 *mco6::HIS3* | This study |
| *mdm10-27* |  | YPH499 *mdm10::ADE2 pFL39-Mdm10-ts27(TRP1)* | This study |
| *mdm10-27 mco6Δ* |  | YPH499 *mdm10::ADE2 pFL39-Mdm10-ts27(TRP1) mco6::HIS3* | This study |

**Table S2:**
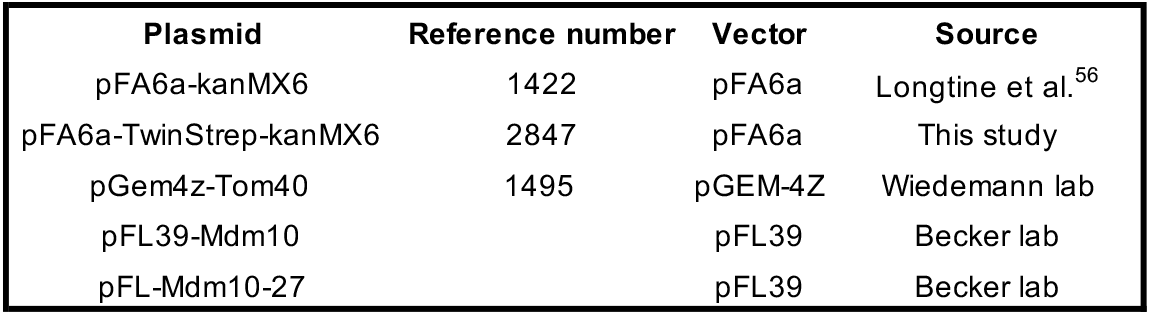
List of plasmids used in this study.

**Table S3:**
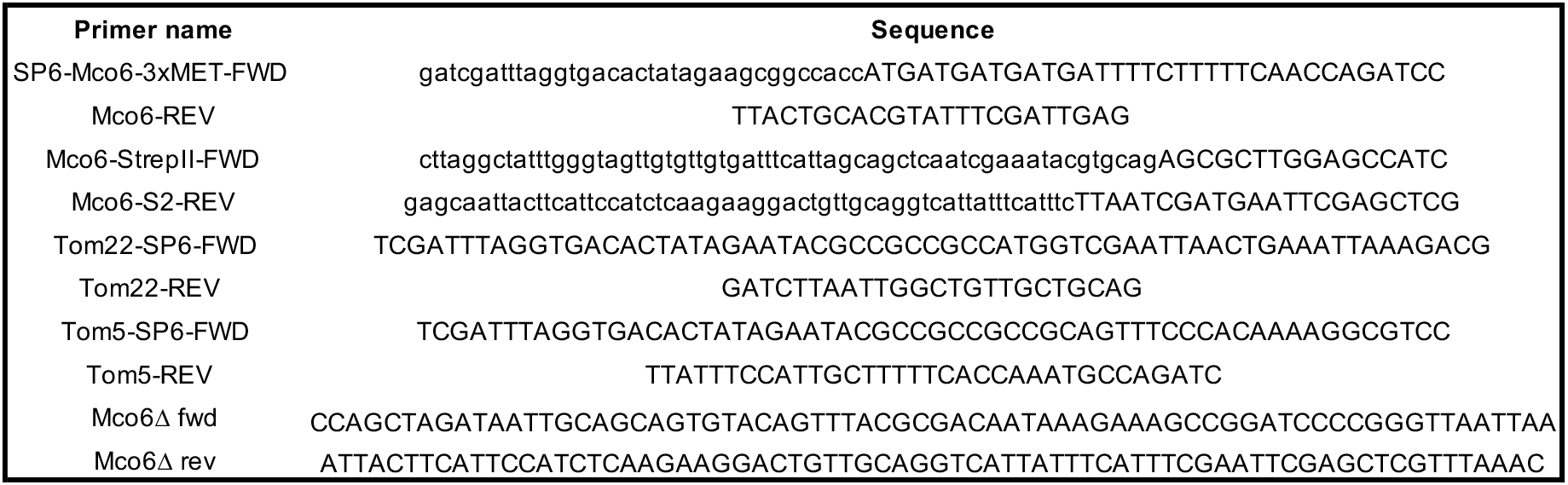
List of primers used in this study.

